# ArthroVerse: mapping protein family diversity across arthropod-associated microbiomes

**DOI:** 10.64898/2026.07.11.737898

**Authors:** Iro N. Chasapi, Eleni Aplakidou, Maria N. Chasapi, Efthimia Lamari, Alexandros Galaras, Sophia Diplari, Ioannis Iliopoulos, Ioannis Z. Emiris, Ilias Georgakopoulos-Soares, Solenn Patalano, Dimitrios J. Stravopodis, Evangelos Karatzas, Fotis A. Baltoumas, Nikos C. Kyrpides, Georgios A. Pavlopoulos

**Author notes:** The first two authors should be regarded as Joint First Authors.

## Abstract

Metagenomic studies of arthropod-associated microbiomes have generated vast amounts of sequence data, yet the functional and structural organization of these proteins remains largely unexplored. Here, we present ArthroVerse, the first comprehensive database of protein families derived from arthropod-associated metagenomes. Non-redundant protein families were generated after rigorous filtering, deduplication, and clustering. The protein families were further annotated with microbial taxonomy, host associations, protein structural information, and Carbohydrate-active enzymes (CAZyme) predictions. The resulting dataset integrates both metagenomic and reference genome-derived proteins, enabling systematic exploration of functional diversity, evolutionary relationships, and host-microbe interactions in insect microbiomes. ArthroVerse provides a valuable resource for the study of microbial ecology and arthropod physiology, offering unprecedented insight into the protein landscape of insect-associated microbial communities.

## Introduction

Arthropods comprise more than 80% of animal species and represent the most diverse and widely distributed phylum on Earth, including insects, arachnids, crustaceans, and myriapods. Their remarkable adaptability to evolutionary pressures, large population sizes, and ecological plasticity have allowed them to diversify and specialize across a range of ecological habitats, ultimately occupying nearly all terrestrial and aquatic ecosystems. Arthropods play fundamental roles in maintaining ecosystem equilibrium through pollination, decomposition, nutrient cycling, and pest regulation (1). At the same time, numerous species, including mosquitoes and flies, serve as vectors of human and animal pathogens, while others act as major agricultural pests (2) or invasive species that threaten biodiversity (3). Beyond their ecological significance, arthropods provide important economic and biotechnological benefits. They contribute to food production through products such as honey and royal jelly produced by the honeybee (Apis mellifera), and many species are increasingly recognized as sustainable protein sources for human consumption and livestock feed (4). In addition, specialized insects produce valuable biomaterials, including silk and other bioactive compounds with applications in medicine, pharmaceuticals, and biomaterials (5).

Arthropod physiology is largely shaped by interactions with their associated microbiomes, which influence host biology, including nutrition, digestion, immunity, development, reproduction, and environmental adaptation (6–8). Recent advances in shotgun metagenomic and 16S rRNA sequencing have revealed complex and diverse microbial communities associated with arthropods, including bacteria, archaea, fungi, and viruses (9–11). Specifically, insect guts are dominated by bacterial phyla such as Proteobacteria, Bacteroidetes, and Firmicutes, whereas specialized diets and nutritional conditions require distinct microbial taxa and functions (12). For instance, termites that feed on wood and dendrites harbor methanogenic archaea or fungi that produce enzymes that degrade lignocellulose into glucose (11).

Mutualistic relationships with bacteria are a hallmark of arthropod-microbiome interactions, with symbiotic lineages having coevolved with their insect hosts over millions of years. Symbionts may be acquired horizontally from the environment or vertically transmitted across host generations. Obligate endosymbionts reside in specialized gut compartments called bacteriocytes (13) and provide essential nutrients absent from nutritionally restricted diets, while facultative symbionts confer context-dependent benefits, including defense against natural enemies and environmental stressors (14).

Despite growing interest in arthropod-related microbiomes, the field lacks specialized resources. Existing databases such as the i5k initiative (15), FlyBase (16), Arachnoserver (17), and the Hymenoptera Genome Database (18) primarily focus on arthropod genomes and do not include their associated microbial communities. BeeBiome represents a notable exception, providing an actively maintained portal dedicated to microbiomes of Apidae species (19). Broader platforms such as IMG/M (20) and MGnify (21) contain extensive metagenomic datasets, but are not specifically tailored to arthropod microbiomes, limiting their utility for studies of arthropod-associated microbiomes. To our knowledge, there is currently no dedicated resource for the systematic exploration of proteins encoded by arthropod-associated microorganisms.

Here, we present ArthroVerse, a comprehensive database of protein families derived from arthropod-associated metagenomes worldwide. ArthroVerse integrates Pfam and CAZyme annotations, microbial taxonomy, host associations, and protein structural predictions to provide a unique framework for investigating microbial function in arthropods. Building on our previous efforts to characterize protein families (22, 23) in specific habitats through the EnvoFams portal (https://envofams.org) (24–28), ArthroVerse enables researchers to link arthropod hosts with their microbial communities at the functional level.

## Materials and Methods

### Data collection and filtering

All publicly available arthropod-associated metagenomes, metatranscriptomes, and reference genomes from the Integrated Microbial Genomes & Microbiomes (IMG/M) database (29), hosted by the Joint Genome Institute (JGI), were collected for the analysis in September 2025. The collected dataset comprises 445 taxon IDs, including 337 metagenomes and 108 metatranscriptomes. Additionally, it contains 3,694 reference genomes, consisting of 2,237 bacterial, 1,409 viral, 44 archaeal, and 4 eukaryotic datasets.

The metagenome and metatranscriptome datasets (environmental samples) were further filtered by matching IMG/M taxon IDs to their corresponding GOLD Sequence Project IDs (30), since multiple IMG/M taxon IDs can map to the same GOLD Sequence Project. To ensure a single representative sample per GOLD study, only one sample per project was retained, selecting the sample with the highest number of predicted proteins. This resulted in a dataset comprising 74,894,168 proteins derived from 368 unique environmental samples. Several stringent filtering steps were applied to ensure the reliability and quality of the predicted proteins (Figure 1). Sequences located within 10 nucleotides of scaffold edges were excluded to minimize truncation artifacts. Additionally, only scaffolds longer than 500 nucleotides were retained to avoid fragmented assemblies. Low-complexity regions were masked using tantan (version 26) (31). Protein-coding sequences shorter than 35 amino acids were removed to reduce the inclusion of non-functional predictions. Finally, MMseqs2 Linclust (32, 33) was applied with a 100% sequence identity threshold and 100% coverage to remove exact duplicates and generate a non-redundant protein dataset without loss of biological information. After filtering and deduplication, the final dataset comprised 53,755,883 non-redundant protein sequences derived from 23,854,349 scaffolds and 340 environmental samples.

**Figure 1.**
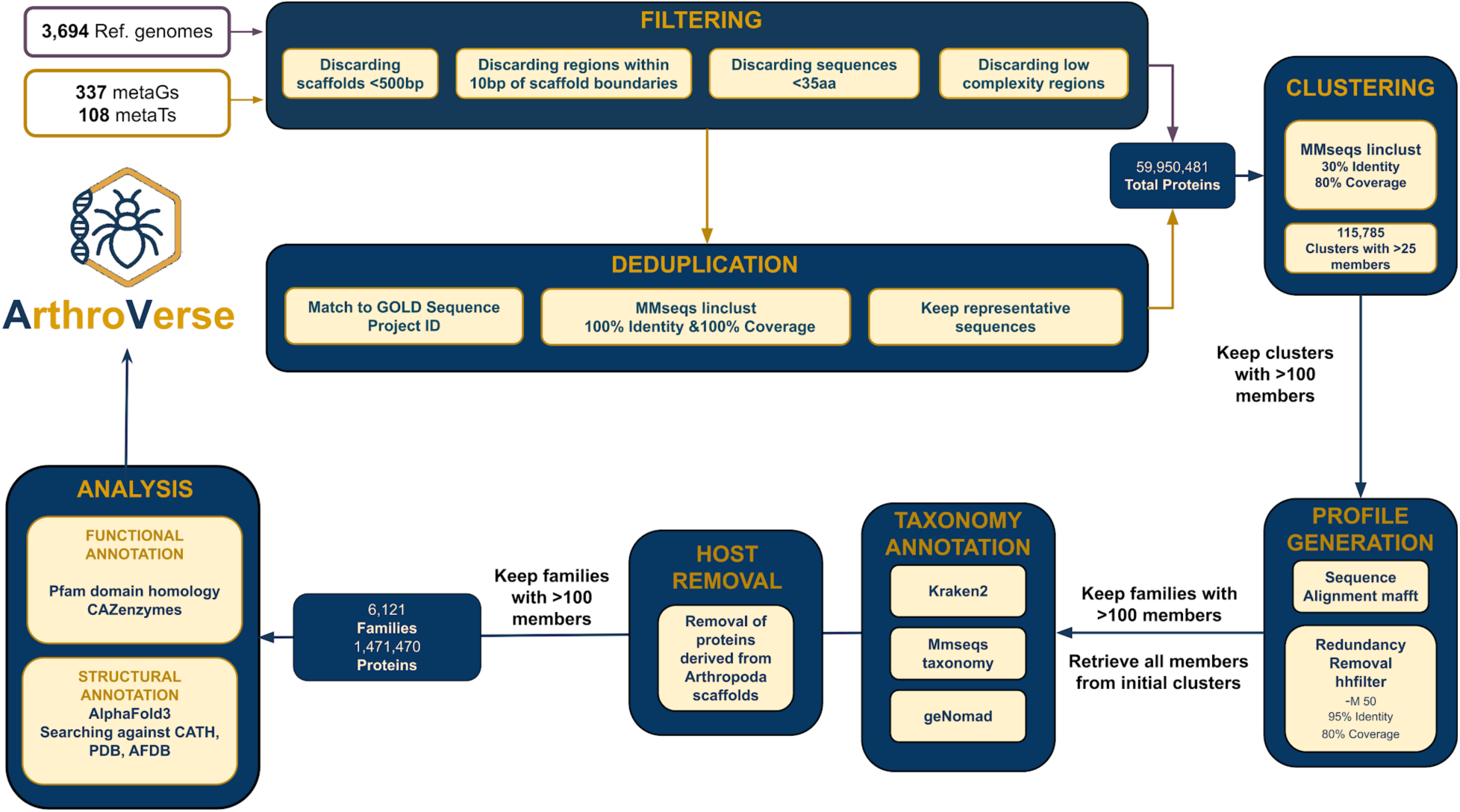
Pipeline overview.

Proteins from reference genomes were filtered using the same quality criteria, including a minimum protein length of 35 amino acids and the removal of low-complexity regions using tantan (version 26). A total of 6,194,598 proteins were retained from 3,694 reference genomes. Filtered proteins from environmental samples and reference genomes were combined into a single dataset of 59,950,481 proteins representing the arthropod microbiome.

### Protein family generation

For protein family generation, MMseqs2 Linclust was applied again, with parameters set to 30% identity and 80% coverage. MAFFT (34) was utilized to align 14,306 selected protein families with 100 or more members. After aligning the sequences, hhfilter (35) was used to remove redundancy, with parameters set to a 95% identity threshold and 80% coverage. The number of families after hhfilter that have 100 or more members is 6,244. The members from the initial clusters are then retrieved. Finally, trimming was applied in order to remove poorly aligned, low-quality, or uninformative regions from a multiple sequence alignment. These regions often contain many gaps, high variability, or low conservation and usually do not reflect true evolutionary or structural signals. The final dataset consists of 6,244 families, yielding 804,555 scaffolds and 1,510,593 proteins, of which 1,173,626 are derived from metagenomes (77,7%) and 336,967 from reference genomes (22,3%).

### Taxonomy classification

The taxonomic analysis was restricted to protein families containing more than 100 members to maintain statistical reliability. Scaffold sequences associated with these protein families were obtained from the IMG/M database. Taxonomic classification was initially conducted using Kraken2 (36). To improve the identification of eukaryotic proteins and limit erroneous assignments, the classifications were refined using MMseqs taxonomy (37). In addition, GeNomad (38) was applied to detect viral and plasmid sequences, ensuring comprehensive identification of mobile genetic elements. Together, these analyses identified 689,291 scaffolds from bacteria (85.6%), 4,031 from archaea (0.5%), 383 from viruses (0.04%), 15,485 from eukarya (1.9%), and 95,365 (11.8%) remaining unclassified.

### Host decontamination

To remove host contamination from our samples, all scaffolds classified as Arthropoda were removed. Following this removal, protein family sizes were re-evaluated, and families with fewer than 100 members were excluded. After this final filtering step, the dataset comprised 6,121 families, totaling 1,471,470 proteins, derived from 778,149 scaffolds, of which 1,136,312 (77.2%) were from metagenomes and 335,158 (22.8 %) from reference genomes.

### Host Distribution Analysis

As part of the functional analysis, the distribution and sharing of protein families across different arthropod orders (e.g., Coleoptera) were examined. Information on arthropod host taxa associated with reference genomes and metagenomes was retrieved from the Integrated Microbial Genomes (IMG) database. Host organisms were classified at the arthropod order level, and protein families were grouped according to this taxonomic assignment. This classification strategy was adopted to preserve ecologically and biologically relevant information while enabling comparative analyses across major arthropod lineages. Organizing host data at the order level facilitated the assessment of protein family presence and overlap among arthropod groups, while minimizing the loss of taxonomic resolution associated with broader classifications. Several protein families exhibited more generic functional roles, as they were shared across multiple arthropod orders.

### Structural model prediction and annotation

AlphaFold3 (39) was used to generate structural models for each protein family. A FASTA file containing all aligned and trimmed member sequences was created for each family and was then converted into an AlphaFold3-compatible JSON input. The representative sequence in each file was treated as the query sequence, while the full FASTA alignment was provided as the unpaired MSA. No paired MSAs or structural templates were used. AlphaFold3 inference was performed using the official model parameters, and one structural model was generated per family. Confidence metrics, including mean pLDDT and pTM, were extracted from the AlphaFold3 output and used to assess model quality. The resulting models were compared to experimentally validated structures from CATH (v4.4, February 2023) (40) and PDB (2024 release) (41), as well as both experimental and predicted structures in AlphaFoldDB (42), using Foldseek (43).

### Functional annotation

The generated protein families were functionally annotated using Pfam (44), a secondary database integrated within InterPro (45). As of December 2024 (Pfam version 38.0), Pfam comprised 25,545 protein families and 796 clans. Functional annotation was performed using hmmsearch from HMMER (version 3.3.2) (46). Of the 6,121 protein families generated, 5,847 had at least one match in Pfam, while the remaining 274 appear to contain potentially novel sequences.

### CAZyme annotation

Representative sequences from each protein family were screened against the dbCAN v12 database using dbCAN3 (47) with both HMMER-based (48) and DIAMOND-based (49) annotation pipelines. Only CAZyme predictions supported by both methods were retained as high-confidence annotations, in accordance with the recommended dbCAN3 workflow (47). To further explore CAZyme categories across microbial families, scaffold taxonomy was manually curated and organized into aerobic and anaerobic groups.

### Database implementation and structure

ArthroVerse is an ASP.NET Core MVC application that uses DuckDB (50) for data management, written primarily in C# with Razor Pages for the interface and CSS/JavaScript for styling. The architecture follows a layered design: C# model classes define core entities with validation; service classes handle data access and processing via DuckDB; controllers manage user interactions and route results; and Razor views render the output. Factory classes centralize object creation to reduce component coupling. The system integrates several external tools for sequence analysis and visualization: DIAMOND for sequence alignment, HMMER for HMM-based homology searches, and ReSeek (51) for additional sequence search. Visualization is supported by the nextProt Feature Viewer (annotated sequence features) (52), MSAViewer (multiple sequence alignments), SkyLign (53) (sequence logos for HMMs and residue frequencies), Mol* (54) (3D molecular structure visualization), and OpenStreetMap (geospatial data).

## Results

### Database components

ArthroVerse catalogs 6,121 non-redundant protein families derived from 1,471,470 proteins associated with 126 insect hosts. The database was constructed using 337 metagenomes, 108 metatranscriptomes, and 3,694 reference genomes from IMG/JGI, ultimately including 126 insect hosts (*Figure 1*). Only protein families comprising at least 100 members were further characterized through functional and structural annotations. ArthroVerse enables exploration of protein families through MSAs and HMMs, AlphaFold3-derived structural models, and taxonomic and host-ecosystem metadata.

### The ArthroVerse database and functionality

ArthroVerse is organized around a top navigation bar that provides access to all major database functionalities, including browsing protein families, scaffolds, samples, host associations, and geographical sample distributions, as well as sequence-search tools and summary statistics (*Figure 2A*).

**Figure 2.**
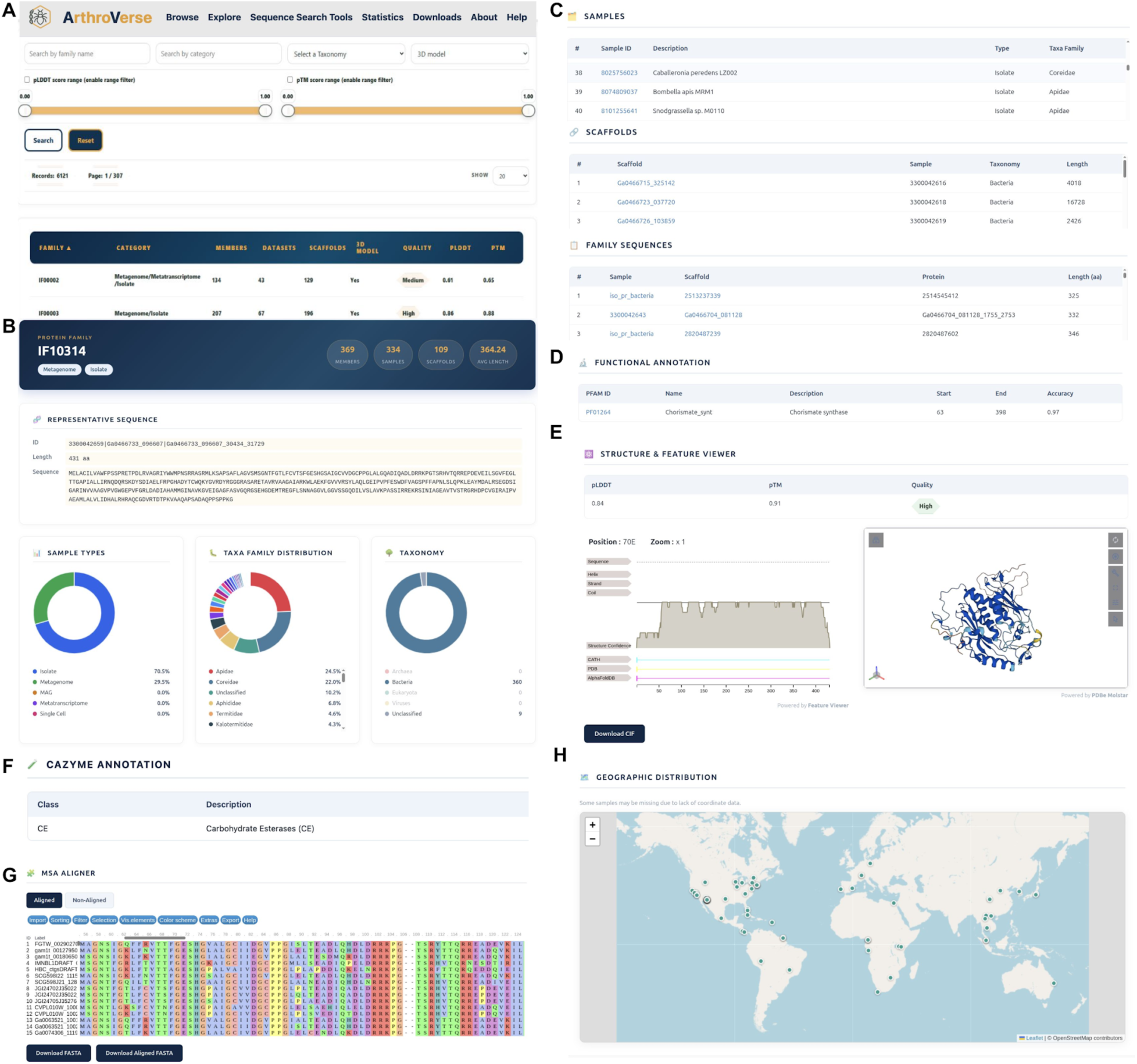
ArthroVerse interface. (A) The Browse Families page offers free-text search and filtering by sequencing category, taxonomy, and structural prediction metrics. (B) The protein family page displays the protein identifier, the number of members, associated datasets and scaffolds, the average sequence length, and the representative amino acid sequence. Interactive pie charts show the distribution of sequencing categories, host taxonomy, and microbial taxonomy. (C) Tables display the samples, scaffolds, and protein sequences linked to the family. (D) Additional tables report Pfam functional domains and top structural matches. (E) Interactive viewers include a sequence logo from the profile HMM with export options; a linear feature map of topology, secondary structure, conservation, and domain annotations; an embedded AlphaFold3 3D model. (F) Tables that display DIAMOND and HMMER similarity of a protein family to CAZymes (if applicable). (G) MSA visualization with customization and FASTA export. (H) Geographical distribution of sampling sites associated with the specific protein family.

Protein families can be explored through both simple and advanced query interfaces. Families can be retrieved through free-text searches using family identifiers and further filtered according to sequencing category (metagenome-only, metatranscriptome-only, mixed, or all families), microbial taxonomy, host association, and structural prediction quality (*Figure 2A*). Additional filtering based on AlphaFold confidence metrics, including pLDDT and pTM scores, is available through interactive range sliders.

The page for an individual protein family displays its unique identifier along with key metadata, including the number of family members, associated datasets and scaffolds, and the average sequence length (*Figure 2B*). Metadata for the representative sequence are also provided, with all identifiers linked directly to IMG/M. (*Figure 2B*). Associated datasets and scaffolds are presented in interactive tables accompanied by their corresponding metadata (*Figure 2B*). Pie charts further illustrate the distributions of sequencing types, arthropod hosts, and microbial taxa associated with the specific family (*Figure 2B*). Additional tables present detailed information for all samples, scaffolds, and family sequences retrieved from IMG and linked to the selected protein family (*Figure 2C*).

Functional annotation of each protein family is provided through Pfam predictions, including domain positions and scores (*Figure 2D*). Structural annotation reports the top five matches against the PDB, CATH, and AlphaFoldDB for the representative sequence, along with TM-scores and alignment coverage to facilitate assessment of structural similarity (*Figure 2E*). Moreover, for protein families identified as CAZymes, ArthroVerse displays an additional table summarizing DIAMOND and HMMER similarity scores against reference CAZyme entries (*Figure 2F*).

Several interactive tools are available for exploring protein family sequences. Multiple sequence alignments (MSAs) can be visualized in either full mode, displaying all family members, or seed mode (*Figure 2G*). The alignment viewer supports custom coloring schemes, identity and occupancy thresholds, and motif or regular-expression searches. An adjacent sequence logo viewer renders the family’s profile HMM as an interactive plot of residue probabilities (each position being clickable to reveal detailed scores). Both the MSA sequences and the underlying HMM can be exported in FASTA, HMMER, or HH-suite format accordingly. Per-residue annotations, including transmembrane topology, secondary structure elements, conservation profiles, and domain predictions (e.g., Pfam), are mapped onto the representative sequence using a linear feature viewer (*Figure 2E*). When available, AlphaFold models are displayed as interactive, rotatable three-dimensional structures and can be downloaded in CIF format (*Figure 2E*). Finally, an interactive map depicts the geographic distribution of sampling locations when coordinate information is available in IMG/M (*Figure 2H*).

A separate section is dedicated to advanced search options focusing on host and microbial taxa, as well as their symbiotic relationships. To support these features, host metadata were manually curated and standardized, harmonizing host names and taxonomic annotations across all datasets. In parallel, our custom taxonomic assignment pipeline integrates multiple complementary classification tools, ultimately achieving greater taxonomic depth and sensitivity. Unlike generic microbiome resources such as Integrated Microbial Genomes & Microbiomes (IMG/M) and MGnify, Arthroverse is the first platform to directly integrate arthropod host identity, microbial taxonomy, and functional annotations, enabling comprehensive exploration of host–microbe–function relationships.

Users can directly query arthropod hosts of interest, including species-level searches. Selecting a host retrieves all protein families exclusively associated with that host, facilitating the identification of host-specific microbial taxa and their functional repertoires (*Figure 3A*). Likewise, microbial taxa can be queried at species-level resolution to retrieve their associated protein families (*Figure 3B*). To further explore host–microbe interactions, Arthroverse integrates host, microbial, and functional information, enabling visualization of the microbial species and their associated protein families detected for each arthropod host (*Figure 3C*). These associations are presented as an interactive network in which the selected host is represented as the central node, connected to its associated microbial species, with edges color-coded by microbial taxonomic family, facilitating the identification of taxonomic patterns within host-associated communities (*Figure 3D*).

**Figure 3.**
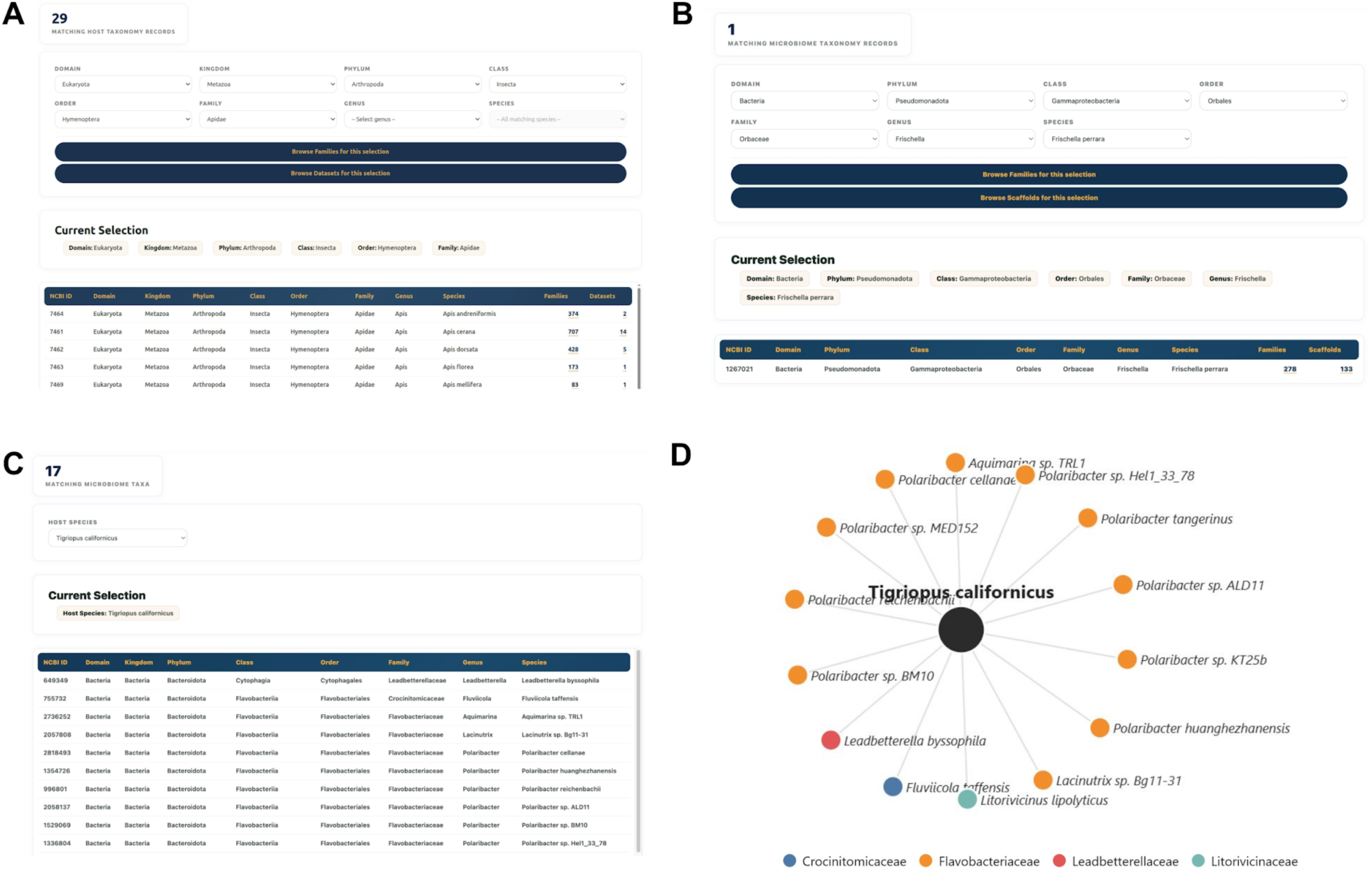
ArthroVerse’s Host–Microbe–Function exploration axis. (A) Explore by Host allows users to select a host at any taxonomic level and view the associated microbial protein families. (B) Explore by Microbiome enables users to select microbial species and view the protein families they encode. (C) The Symbiosis page displays the taxonomic classification (Domain, Kingdom, Phylum, Class, Order, Family, Genus, and Species) of microorganisms associated with the selected host. (D) The Host–Microbiome Network visualizes the microorganisms associated with the selected host. The central node represents the host, while the surrounding nodes represent microbial taxa, color-coded by taxonomic family.

A dedicated section of sequence search tools is available for users to query their sequences of interest against the ArthoVerse database. Sequence similarity and HMM-based searches are supported by DIAMOND (49) and HMMER (48), respectively, and users can customize parameters such as score cutoffs, substitution matrices, and gap penalties. PROSITE-style motif queries or arbitrary regular expressions can be searched through a dedicated interface, enabling the detection of conserved sequence features across protein families. Users can also apply Reseek (51) with adjustable sensitivity modes, E-value thresholds, and query coverage to perform structural similarity. All results tables are downloadable for further processing.

### Data statistical analysis

The statistical metrics of the final dataset highlight the diversity, scale, and quality of the generated protein families from arthropod-associated microbiomes. The distribution of protein families across datasets showed that most protein families were niche-specific. The vast majority of families appeared in only 51-100 datasets, while a significant portion was found in 50 or fewer (*Figure 4A*). Only a small fraction of protein families was found in >500 datasets, suggesting microbial protein specificity across diverse arthropod hosts and environments. Moreover, most families included members encoded by 101-200 scaffolds (*Figure 4B*). We observed that the majority of protein clusters were small, with more than 3,000 families containing between 101 and 190 members (*Figure 4C*).

**Figure 4.**
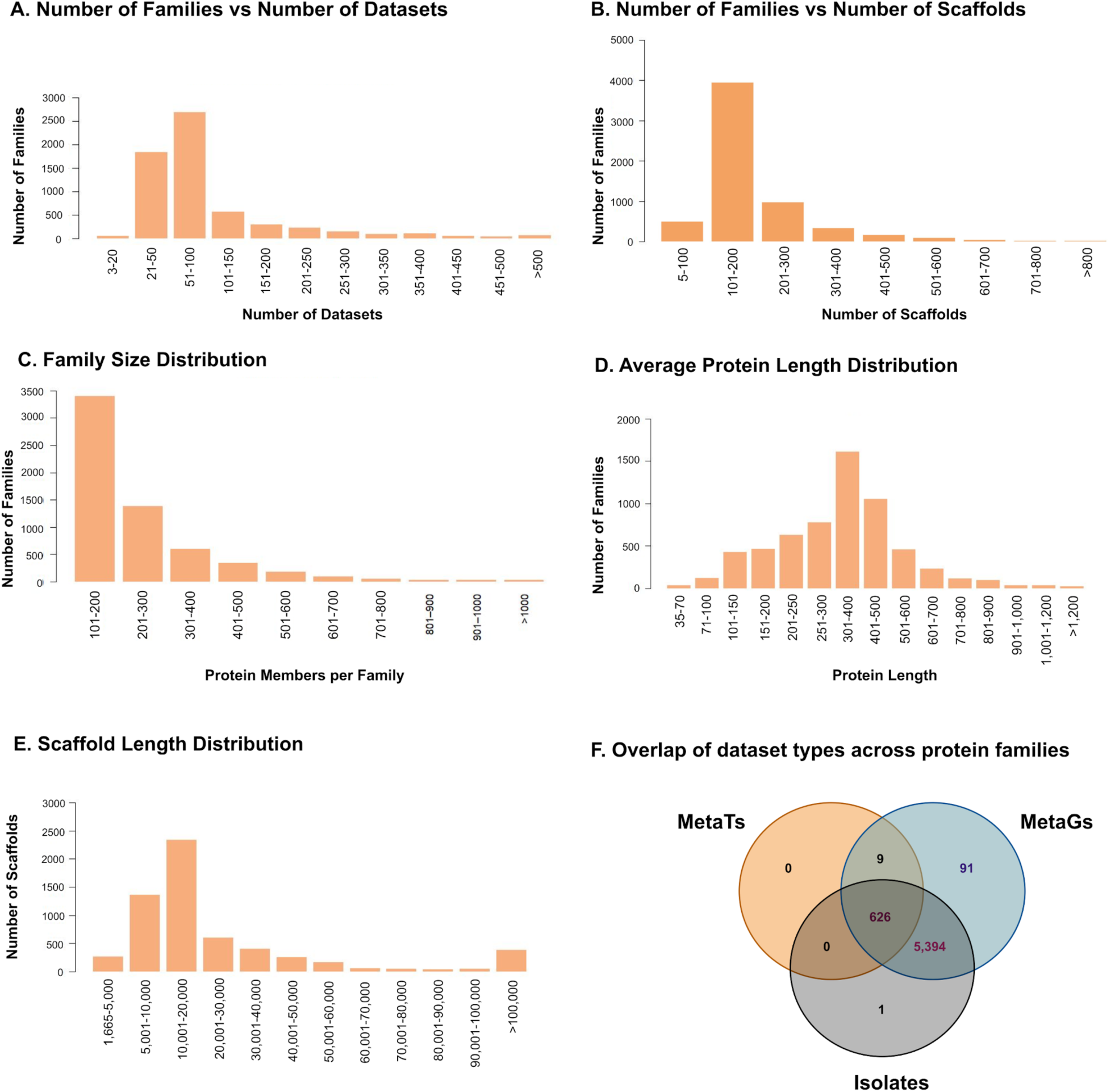
Dataset statistics. (A) Number of Families vs Number of Datasets. Number of families relative to the number of datasets, showing that most families comprise 50–150 datasets. **(B) Number of Families vs Number of Scaffolds.** Number of families relative to the number of scaffolds, highlighting that most families derived from 100-200 scaffolds. (**C) Family Size Distribution.** Histogram showing the number of proteins per family, highlighting that most families contain 100–300 members. **(D) Average Protein Length Distribution.** Distribution of the average length of protein members within each family. Based on the filtering criteria, the minimum protein length is 35 amino acids (aa), with most falling between 250 and 500 aa, consistent with typical full-length proteins. **(E) Scaffold Length Distribution.** Distribution across families shows a minimum scaffold length of 1,665 bp and a predominant range of 10,000–20,000 bp. **(F) Overlap of dataset types across protein families**. Venn diagram illustrating the distribution of dataset types, Metatranscriptomes (MetaTs), Metagenomes (MetaGs), and Isolates, represented within the identified families. A stark majority of families are shared exclusively between MetaGs and Isolates, while 626 families are common to all three dataset types

Average protein lengths followed a bell-shaped curve peaking between 301 and 400 amino acids (*Figure 4D*), as expected for microbial proteins, suggesting that our pipeline successfully recovered complete coding sequences. Most scaffolds were around 5,001–15,000 bp in length, while we detected a subset exceeding 100,000 and 200,000 bp (*Figure 4E*). These long scaffolds indicate that most identified protein families are encoded from genomic fragments large enough to contain multiple complete genes, ultimately enhancing precise functional annotation and strengthening taxonomic assignment.

Most protein families were derived from reference genomes (6,021 unique families), followed by metagenomes (MetaGs; 1,352 unique families), whereas metatranscriptomes (MetaTs) contributed no unique families. Most shared families were detected between reference genomes and MetaGs, including 5,394 families overlapping exclusively between these datasets and 626 families shared across all three dataset types (*Figure 4F*). To structurally annotate the protein families, we performed FoldSeek on three major structural databases: AlphaFold (AFDB), CATH, and PDB. A vast majority (>90%) of families showed high structural homology, and 4,918 families predicted to have high-quality structures were shared across all three databases. This level of overlap confirms that the dataset contains well-folded, biologically relevant protein structures with only a very small fraction of families falling into the “Low Confidence” category.

## Case Study 1. Integrative host, taxonomic, and structural analyses uncover termite-associated CAZyme families

CAZymes are responsible for the synthesis, degradation, and recognition of complex carbohydrates and play a central role in arthropods’ ability to exploit plant-derived polysaccharides. CAZymes are classified into six major categories: (i) glycoside hydrolases (GHs), which hydrolyze glycosidic bonds; (ii) glycosyltransferases (GTs), which catalyze the formation of glycosidic bonds; (iii) polysaccharide lyases (PLs), which cleave uronic acid-containing polysaccharides through non-hydrolytic mechanisms; (iv) carbohydrate esterases (CEs), which remove acyl groups from carbohydrate esters; (v) auxiliary activities (AAs), which comprise redox enzymes that facilitate the degradation of recalcitrant polysaccharides; and (vi) carbohydrate-binding modules (CBMs), which mediate carbohydrate recognition and binding (55). Because CAZymes are primarily contributed by the gut microbiota, whereas animals produce only a limited subset, a diverse CAZyme repertoire enables hosts to utilize a broader range of dietary substrates (56).

Our repository enables the systematic exploration of CAZyme repertoire in arthropod-associated microbiomes. We identified 139 protein families as *bona-fide* CAZymes, with GHs and GTs constituting the dominant classes (*Figure 5A*). Taxonomic profiling revealed that GHs, GTs, and CEs are primarily associated with anaerobic bacteria, particularly members of Bacteroidota, Spirochaetota, and Clostridia (*Figure 5B*). In contrast, AAs and CBMs are primarily associated with aerobic microorganisms, mainly Proteobacteria, and are entirely absent from Spirochaetota (*Figure 5B*). The observed distribution of AAs is biologically plausible, as many AA enzymes are oxidative and require molecular oxygen for their catalytic activity (55). Interestingly, despite the strong representation of termite-associated microbiomes in our dataset, AA-containing protein families were not detected in termite hosts. This finding is consistent with the largely anaerobic conditions that characterize termite gut ecosystems and suggests that oxidative carbohydrate degradation plays a limited role in these communities. Our results suggest that the CAZyme repertoire is strongly influenced by the host’s living conditions.

**Figure 5.**
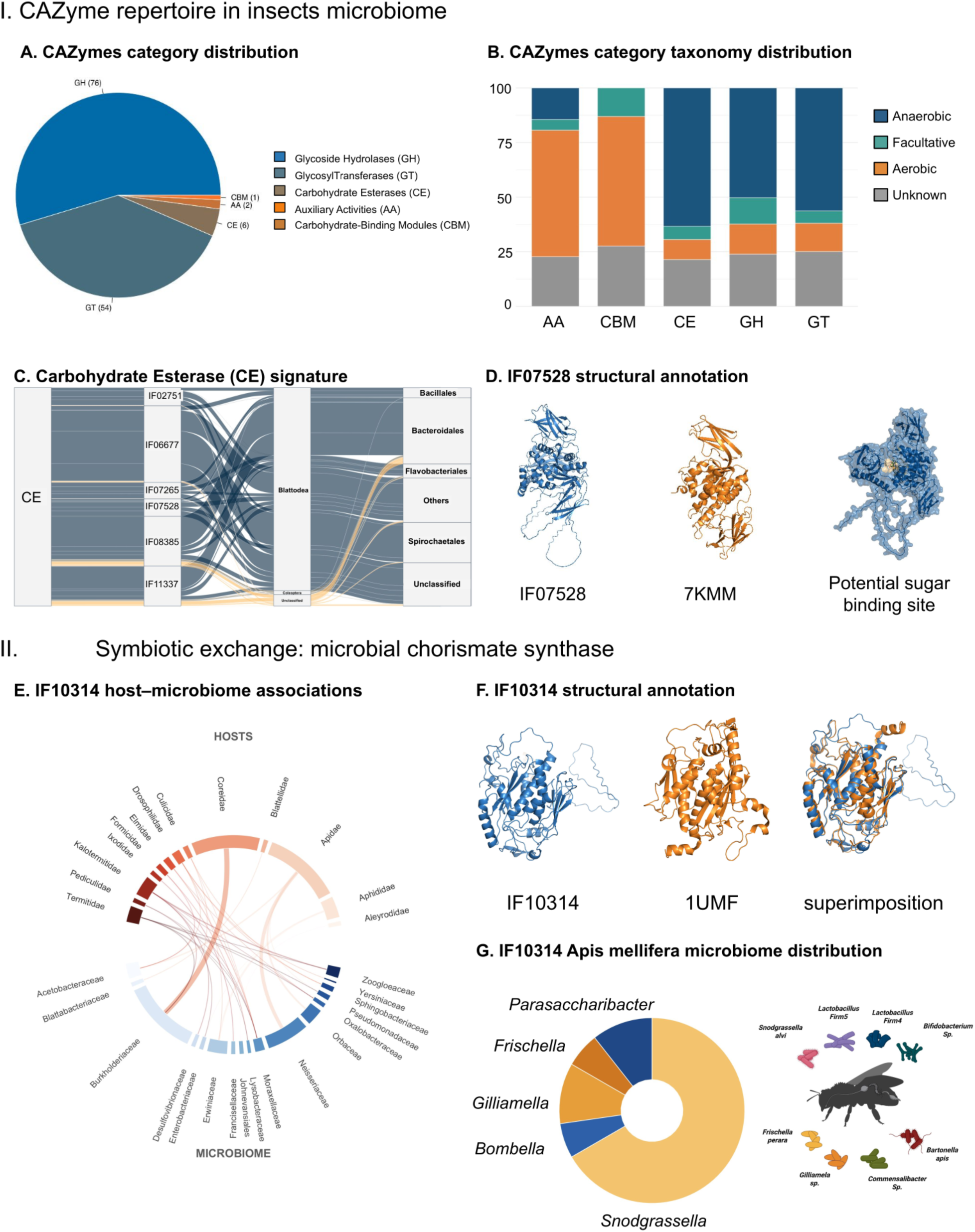
Case studies using ArthroVerse. (I) CAZyme repertoire in insect microbiomes. (A) CAZyme category distribution. Pie chart showing the distribution of CAZyme categories annotated across ArthroVerse families. **(B) CAZyme category taxonomy distribution.** Barplot showing the taxonomic distribution of proteins within families belonging to a specific CAZyme category. Proteins are classified as Aerobic, Anaerobic, Facultative, or Unknown to further explore the prevalence of aerobic and anaerobic organisms within specific categories. Notably, Carbohydrate Esterases (CE), Glycoside Hydrolases (GH), and GlycosylTransferases (GT) are dominated by anaerobic organisms. **(C) Carbohydrate Esterase (CE) signature**. To further explore the taxonomic signatures of families in the CE category, the origins of these families in hosts and the microbiome were determined. The flowchart highlights specific links between hosts and their microbiomes that contribute to the production of these proteins. **(D) IF07528 structural annotation.** A specific family within the CE category was selected. The IF07528 structure was displayed and superimposed against its top PDB hit (7KMM), a novel carbohydrate acetylesterase from Xanthomonas citri. **(II) Symbiotic exchange: microbial chorismate synthase. (E) IF10314 host-microbiome associations.** A chord diagram represents the microbial taxonomy origin of the proteins and their associated hosts in the IF10314 family. **(F) IF10314 structural annotation.** Structural model representation of IF10314, its top PDB structural hit 1UMF (monomer; crystal structure of chorismate synthase), and their superimposition. **(G) IF10314 distribution in the Apis mellifera microbiome.** (Left) Donut plot representing the taxonomic distribution of the IF10314 family proteins. (Right) Gut microbiome taxonomy of Apis mellifera.

Next, we investigated the six protein families classified as carbohydrate esterases (CEs) within the arthropod-associated microbial proteome. Remarkably, 93% of all CE-containing proteins were associated with Blattodea (termites), whereas only a small fraction of IF08385 and IF11337 originated from Coleoptera or unclassified hosts (*Figure 5C*). Taxonomic profiling further revealed that these CE families are predominantly encoded by members of Bacteroides, Parabacteroides, and the spirochaetes Treponema and Breznakiella (*Figure 5C*). These taxa constitute the canonical lignocellulose-degrading community of the termite hindgut and play central roles in the deconstruction of plant biomass (57–59). Carbohydrate esterases remove acetyl and other ester groups from hemicellulose, debranching the polymer and making its backbone accessible to glycoside hydrolases (55, 60). Our findings support that the strong enrichment of CE families in termite-associated microbiomes reflects the specialized adaptation of the termite gut microbiota for efficient lignocellulose degradation.

To further investigate the termite-enriched CE family IF07528, we examined its predicted three-dimensional structure (*Figure 5D*). Structural comparison revealed a high degree of similarity between family IF07528 and the experimentally determined structure 7KMM, which corresponds to a carbohydrate acetylesterase. Superposition of the two proteins revealed a conserved surface pocket predicted to function as a sugar-binding site (*Figure 5D*), validating the family’s classification as CE. Collectively, our case study illustrates how ArthroVerse can integrate host specificity, taxonomic context, and structural annotation to expand our understanding of arthropod-associated microbiomes.

## Case Study 2. Host-specific microbial partners for chorismate synthesis in arthropod microbiomes

The Shikimate pathway is an essential seven-step metabolic process that converts phosphoenolpyruvate and erythrose 4-phosphate into chorismate, the precursor of the aromatic amino acids phenylalanine, tryptophan, and tyrosine, as well as numerous secondary metabolites (61). These amino acids are critical for protein synthesis and the production of molecules necessary for animal physiology. However, insects, like most animals, lack a complete shikimate pathway and therefore cannot synthesize aromatic amino acids de novo, relying instead on dietary sources or metabolic contributions from their symbiotic microbiota (61, 62).

To investigate the microbial potential for aromatic amino acid biosynthesis, we leveraged ArthroVerse to identify protein families associated with the shikimate pathway. Among these, we identified family IF10314, whose core region is annotated as chorismate synthase (Pfam), an enzyme that catalyzes the final step of the shikimate pathway and is absent from insect genomes. IF10314 was identified in multiple insect hosts, including bees, termites, ants, flies, and cockroaches (*Figure 5E*). Interestingly, we observed that the microbial taxa encoding this family differed markedly among hosts. Thus, although chorismate synthase appears to be a widespread microbial function, its taxonomic carriers are highly host-dependent, suggesting that different arthropod lineages have recruited distinct microbial partners to fulfill a common metabolic requirement.

To further validate the functional assignment of IF10314, we examined its predicted three-dimensional structure. Structural comparison demonstrated a high degree of similarity between IF10314 and the experimentally determined chorismate synthase structure 1UMF (*Figure 5F*). Superimposition of the two structures revealed strong conservation of the overall fold, supporting the annotation of IF10314 as a bona fide chorismate synthase.

Given the strong association of IF10314 with Apidae, we further examined its taxonomic distribution within the honeybee microbiome. Most IF10314 proteins were encoded by *Snodgrassella*, *Gilliamella*, and *Frischella*, the three main symbionts restricted only to bees (63), followed by *Parasaccharibacter* and *Bombella* (*Figure 5G*). The enrichment of IF10314 within these core bee-specific microorganisms suggests that microbial chorismate synthesis may represent an evolutionarily conserved metabolic function embedded within the bee microbiome. This case study highlights the potential role of ArthroVerse in disentangling the crosstalk between protein functions and host-microbe symbiosis.

## Discussion

ArthroVerse catalogs 6,121 protein families derived from 337 metagenomes, 108 metatranscriptomes, and 3,694 reference genomes associated with arthropod microbiomes, enabling the systematic exploration of microbial diversity across 126 arthropod hosts. By integrating protein families with microbial taxonomy, host associations, functional and structural annotations, and sequence analysis tools, ArthroVerse facilitates the investigation of host–microbiome interactions and the discovery of protein functions across diverse arthropod-associated microbial communities.

Built on publicly available sequencing data, ArthroVerse reflects the current state of arthropod microbiome research, with well-studied hosts such as honeybees and termites represented more extensively than other lineages. Continuous sequencing initiatives are expected to broaden this representation. The clustering strategy (30% sequence identity and 80% coverage, followed by HHfilter at 95% identity and 80% coverage) was selected to balance family coherence with biological diversity, although alternative clustering schemes may be explored in future updates. Similarly, our decision to comprehensively annotate families containing at least 100 members prioritizes robust, well-supported protein families, while smaller families represent a valuable resource for expansion as the database grows. To maximize the accuracy of taxonomic annotation, we combined multiple complementary tools (Kraken2, MMseqs taxonomy, and GeNomad), ultimately resulting in taxonomic assignments for 88.2% of scaffolds associated with well-supported protein families. Finally, functional and structural annotation with PFAM and AlphaFold (followed by FoldSeek) enabled a comprehensive characterization of the protein families.

ArthroVerse represents the first protein-family database dedicated to arthropod-associated microbiomes. Beyond cataloging protein families, the database uniquely links microbial proteins to their taxonomic origins, arthropod hosts, and structural annotations, enabling comparative analyses of host-specific microbial functions across diverse arthropod lineages. Moreover, users can directly explore the functional repertoire of microbes in individual arthropod species and examine host-associated microbiomes on a unified platform at an unprecedented taxonomic scale. Furthermore, ArthroVerse includes numerous protein families that currently lack functional annotation (thereby leaving great space for novelty), representing a valuable reservoir of candidate proteins for future biochemical, structural, and evolutionary characterization.

We strongly believe that ArthroVerse sets the foundation for researchers to systematically decode the functional landscape of arthropod-associated microbiomes, opening new avenues for understanding arthropod–microbe interactions, symbiosis, and evolution.

## Data availability

All data generated within ArthoVerse are freely accessible and downloadable. Multiple sequence alignments and protein family profiles can be exported in standard FASTA and HMMER or HH-suite formats, respectively. Three-dimensional structural models are provided in CIF format, while functional annotations and associated metadata can be downloaded as TSV files. The complete collection of raw data and downloadable files is archived at Zenodo (doi:10.5281/zenodo.21296430). ArthoVerse is integrated into the EnvoFams portal (https://envofams.org), a comprehensive platform for exploring and characterizing microbial and viral protein families across diverse ecosystems. The ArthoVerse database can be accessed through https://envofams.org/arthroverse.

## Acknowledgements

This work was supported by computational resources provided by the ELIXIR-GR HYPATIA Cloud infrastructure.

